# Abnormal vasculature reduces overlap between drugs and oxygen in a tumour computational model: implications for therapeutic efficacy

**DOI:** 10.1101/2024.09.27.615320

**Authors:** Romain Enjalbert, Jakub Köry, Timm Krüger, Miguel O. Bernabeu

## Abstract

The tumour microvasculature is abnormal, and as a consequence oxygen and drug transport to the tumour tissue is impaired. The abnormal microvasculature contributes to tumour tissue hypoxia, as well as to varying drug penetration depth in the tumour. Many anti-cancer treatments require the presence of oxygen to be fully efficacious, however the question of how well oxygen concentration overlaps with drug concentration is not elucidated, which could compromise the therapeutic effect of these drugs. In this work we use a computational model of blood flow and oxygen transport, and develop a model for an oxygen-dependent drug, T-DM1, to study the overlap of oxygen and drug concentration in healthy and tumour tissue, where we assume the tumour tissue to compress blood vessels. Our results show that, due to the compressed vessels present in tumours, areas of sufficient oxygen concentration for a drug to function overlap poorly with areas of sufficient drug concentration, covering 28% of the tumour tissue, compared to 82% in healthy tissue. The reduction in drug and oxygen overlap is due to the altered red blood cell dynamics through the abnormal microvasculature, and indicates that drug transport to tumours should not be considered independently of oxygen transport in cases where the drug requires oxygen to function.

## Introduction

The abnormal tumour microenvironment (TME) is a hallmark of cancer [1]. The TME impairs drug delivery to tumour tissue and increases tumour tissue hypoxia, both barriers to the clinical efficacy of drugs [2–4].

Pharmacokinetics is an important aspect of the clinical efficacy of a drug [5, 6]. Drugs are first transported through the blood stream and then extravasate to the tumour tissue [2, 7, 8]. Following extravasation, the advection, diffusion and reaction kinetics of the drug determine its transport to tumour tissue [5, 6]. The TME can act as a barrier to drug delivery to the tissue and therefore to its efficacy [5, 9]. However, the presence of a drug can be insufficient to fully determine a drugs efficacy, as recent studies have shown that the cytotoxicity of some drugs may also be reduced by the absence of oxygen [4, 10–13]. As an example, the efficacy of trastuzumab-emtansine (T-DM1), a drug used to treat HER2+ breast cancer, is reduced under hypoxic conditions due to caveolin-1 translocation [10], making the transport of oxygen to tumour tissue of particular relevance for the efficacy of such drugs.

Oxygen travels bound to red blood cells through microvascular networks prior to being delivered to the tissue [14]. In microvascular networks, red blood cells partition unevenly at microvascular bifurcations [15]. Previous work has shown that in networks with tumour vascular abnormalities [16], such as reduced inter-bifurcation distances [17] and vessel compression [18, 19], the partitioning of red blood cells at bifurcations is abnormal, leading to a high heterogeneity of discharge haematocrit (flowrate fraction of red blood cells in blood) [17–20]. As a consequence of haematocrit heterogeneity, blood vessels are inefficient at delivering oxygen and tumour tissue can be hypoxic [17, 21–24]. In addition, tumour microvascular networks can have avascular areas that lead to hypoxic regions due to the limited diffusion distance of oxygen [25, 26].

In cases where a drug requires oxygen to function, the question of the overlap of drug and oxygen concentration for full therapeutic effect needs to be considered. Since oxygen and drugs are transported via different mechanisms, we hypothesise that, in the abnormal tumour microvasculature, areas of high oxygen concentration can overlap poorly with areas of high drug concentration, and that this hypothesis is relevant given the oxygen requirements of some drugs for them to be efficacious. We investigate this hypothesis in a computational model of a tumour with tumour induced compressed vessels,[27] blood flow [18], oxygen transport [17] and T-DM1 transport [7], where the computational model is used as an exemplar to illustrate how the different mechanisms of transport lead to poor overlap between oxygen and drug concentration in the tissue.

Our results show that, due to the abnormal vascular structures present in tumours (vessel compression in our computational model), areas of sufficient oxygen concentration for a drug to function overlap poorly with areas of sufficient drug concentration. This effect is mostly due to the poor oxygen transport to the tumour tissue, and indicates that drug transport should not be considered independently of oxygen transport in cases where the drug requires oxygen to function. These findings provide a theoretical underpinning for therapeutic approaches aimed at enhancing tumour oxygenation in order to improve anticancer drug efficacy.

## Methods

### Tumour and control models

To test our hypothesis, we generate a tumour model and a control model to compare the two. In both models, the same microvascular network is used, illustrated in Figure 1a, and generated using the open-source Tumorcode software [28]. The code employs a set of rules based on Murray’s law and the properties of venules, arterioles and capillaries to generate a microvascular network on a mesh with properties matching real network sample [29]. The settings we use to generate the network are the default two-dimensional network generator with a single inlet and a single outlet in a 5000 *µm* by 5000 *µm* square domain. We separate the microvascular network into periphery and core vessels, shown in Figure 1a, where core vessels are defined as having both ends of the vessel within 1000 *µm* of the centre of the network [18], the remaining vessels are periphery vessels. In the tumour model, the core vessels are compressed but not the periphery ones, while in the control model none of the vessels are compressed.

**Figure 1.**
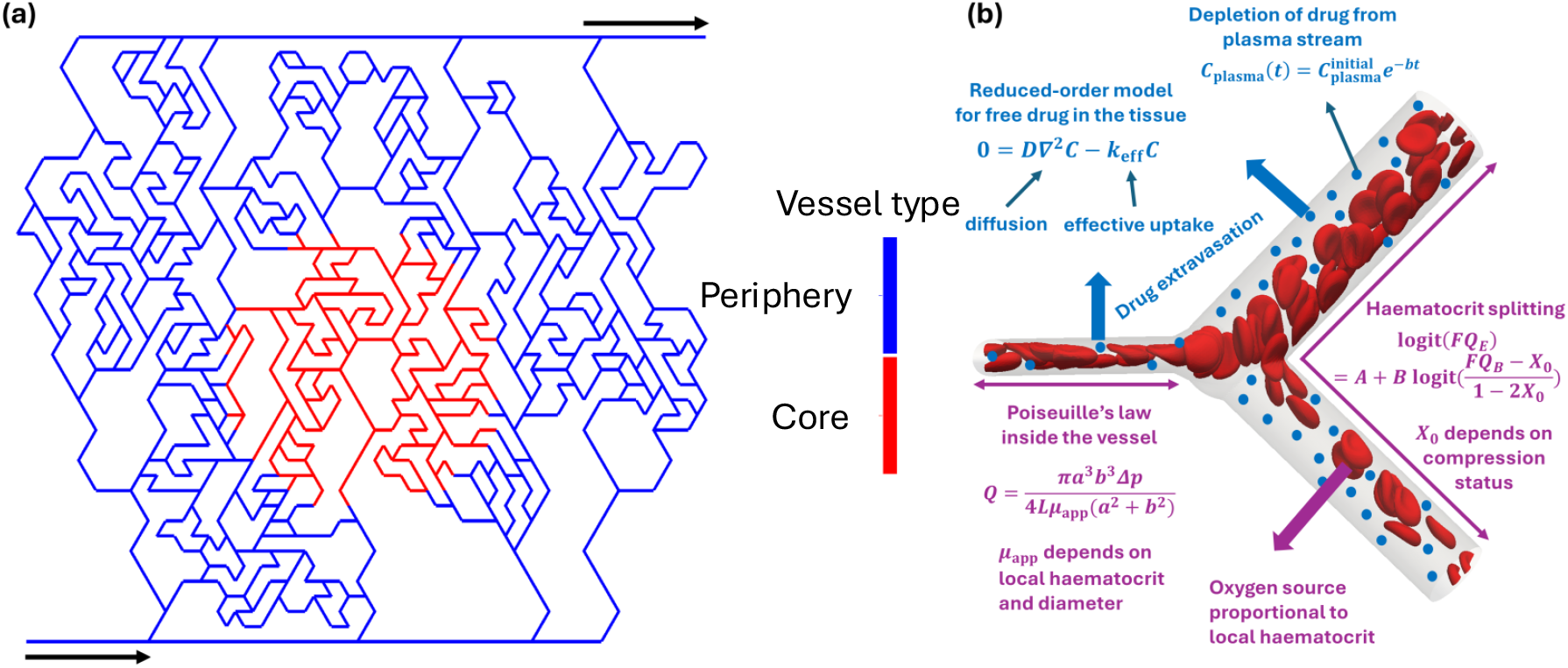
(a) Illustration of microvascular network generated with Tumorcode [28], see methods for more details. The black arrows indicate the inlet, at the bottom left, and the outlet, at the top right. In red are the core vessels (compressed in the tumour model) and in blue are the periphery vessels. (b) Illustration of a diverging vessel bifurcation showing key equations of the presented model at the microscale, the red blood cells are for illustration purposes as they are not individually resolved in our model. The equations and their variables are defined in Methods section and the supplementary material.

### Blood flow

In this work, we treat blood flow as a one-dimensional continuous fluid, imposing Poiseuille’s law at every vessel segment [15]. In addition, we use existing models to implement the Fåhraeus effect, Fåhraeus-Lindqvist effect, and phase separation at bifurcations in our treatment of blood flow [30–32]. In addition, we consider some blood vessels to be compressed, and in those cases treat vessel-cross sections as elliptical (with an aspect ratio of 4.26 [27]) and use a modified phase-separation model for red blood cell partitioning at bifurcations [18]. The networks were chosen to have a single inlet and a single outlet, facilitating the choice of boundary conditions as they do not affect the distribution of discharge haematocrit in the network, therefore an arbitrary pressure value is imposed at the inlet, and a 0 value at the outlet. Unless stated otherwise, the inlet discharge haematocrit to the network is 30% [33]. The supplementary material contains the full details of the blood flow model.

### Oxygen and drug transport

Vascular networks are embedded inside a rectangular domain representing the tissue through which oxygen and T-DM1 diffuse. Both chemical species are uptaken by the tissue and are supplied by vessels acting as line sources with strength proportional to local vessel diameter thus perfusing the tissue.

The oxygen transport problem is governed by the same equation and parameters as described in Supplementary material to [17] and the numerical simulations are performed in Microvessel Chaste [34]. Here, by focussing on the long-time behaviour we adopt a quasi-steady state approximation [35]. Mathematically, the problem thus reduces to a steady reaction-diffusion equation

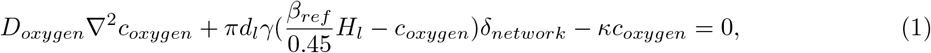

where *D*_*oxygen*_ is the diffusion coefficient for oxygen in the tissue, *c*_*oxygen*_ is the oxygen concentration in the tissue, *γ* is the vessel permeability, *d*_*l*_ is the local vessel diameter, *H*_*l*_ is the local vessel discharge haematocrit, *β*_*ref*_ relates the discharge haematocrit in the vessel to an oxygen concentration in the vessel, *κ* is the rate at which oxygen is consumed by the cells, and *δ*_*newtork*_ is a representation of the vessel network via a collection of Dirac delta function sources [17]. The majority of the governing parameters are taken from [36], while the haematocrit term is taken from the blood flow simulations described in the Blood Flow section.

For the tissue dynamics of T-DM1, one must typically distinguish between three distinct phases of the drug – free, bound and internalized – and model transitions between these phases. Our starting point is the model from [7]. Free drug (concentration *C*) diffuses in the tissue with a diffusion constant *D* and is supplied by the vessels at a rate proportional to the plasma concentration *C*_plasma_ (which decays in time due to depletion of drug from the blood stream, more details to follow in this section). Simultaneously, the free drug binds to receptors embedded in cell membranes (supplied at a fixed concentration *C*_*r*_ for simplicity) to form bound complexes (concentration *B*) with a rate constant *k*_on_ and unbinds from these complexes at a rate *k*_off_. Drug in bound complexes is thus immobilized (cannot further penetrate the tissue) and internalized (concentration *I*) at a rate *k*_int_. As trastuzumab is a monoclonal antibody, we assume no binding to blood cells or plasma proteins occurs (*f*_*free*_ = 1 holds for the free-drug fraction from [5]). The full dimensional model for T-DM1 kinetics then reads

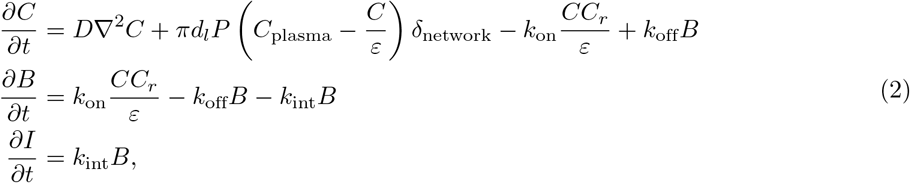

where *d*_*l*_ is the local vessel diameter, *ε* is the effective void fraction, *P* the vessel permeability and *δ*_network_ is a representation of the vessel network via a collection of Dirac delta function sources [7, 17]. Using data from [37] we found (see Supplementary material) that the depletion of T-DM1 from the blood stream can be modelled via an exponential function from data from [37]

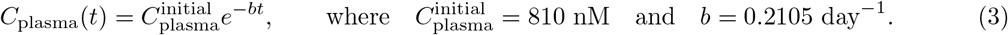

Eq. (2) with plasma concentration depleting according to Eq. (3) features six physical processes - diffusion, extravasation, plasma depletion, binding, unbinding and internalization - and each of these proceeds at an associated timescale. In our network, local vessel diameters *d*_*l*_ and inter-vessel distances *h*_*l*_ are in the ranges 15 − 110 and 100 − 1000 *µ*m, respectively. Using default parameters (as summarized in Table S1), we can then estimate the values and ranges of the 6 timescales present in our simulations as follows:

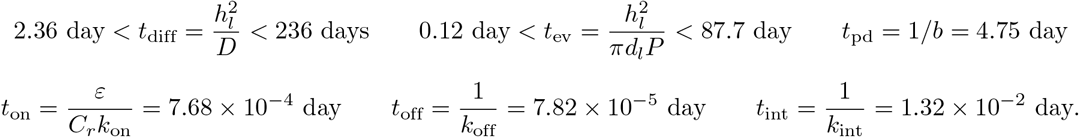

We see that the equilibration of diffusion, extravasation and plasma depletion processes happens much slower (days to months; note that standard T-DM1 treatments are administered in three-week cycles, i.e on the same timescale [37]) than that of kinetics (minutes), i.e.

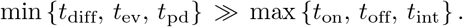

Thus we propose a quasi-static model reduction whereby the equation for *B* is quasi-static (i.e. *∂B/∂t ≈* 0).

This means that *B* quasi-statically follows *C* according to

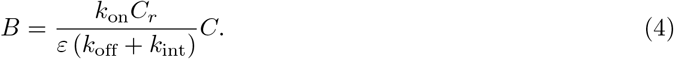

Substituting this result back into the equation for *C*, we obtain

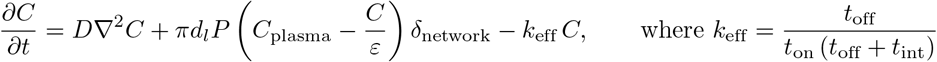

is the effective rate of the drug uptake. For the timescale of the effective uptake process we then have

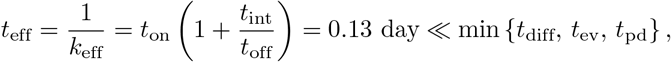

which holds true except for extreme combinations of very large local vessel diameter and very small inter-vessel distance. Thus, the distribution of the free drug on the timescale of interest (large times) follows from a linear quasi-static, which we refer to quasi-steady-state, reaction-diffusion equation

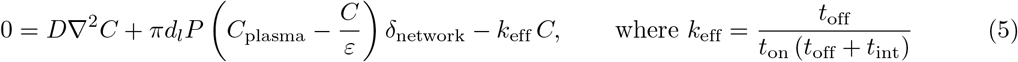

and *C*_plasma_ is given in Eq. (3). The bound-drug concentration *B* then follows from the quasi-steady-state assumption in Eq. (4) and the internalized drug *I* is obtained simply by integrating the last equation in Eq. (2). Detailed derivations pertaining to the proposed model reduction are summarized in Supplementary material. Moreover, when posed on a simple geometry wherein a single cylindrical blood vessel perfuses the surrounding tissue, the reduced-order model is amenable to direct mathematical analysis - we found an exact solution to this problem and confirmed the validity of the model reduction via comparison with the numerical solution of the full problem - see Supplementary material. All dimensional parameters with appropriate units and references are summarized in Supplementary Table S1.

### Processing results

We define hypoxia as an oxygen concentration below 8 mmHg. We choose this value as it is defined as hypoxia in tumours [38], and because the efficacy of T-DM1 is reported to be reduced at that level of oxygenation [10]. Similarly, we determine that there is a sufficient concentration of T-DM1 when it is above the IC_50_ (half maximum inhibitory concentration) reported in the literature, 2.9 nmol/L or 6.8 nmol/L, depending on the tumour cell line [10]. Therefore, there are two conditions required for optimal drug effect, which are a sufficiently high drug concentration, above the IC_50_, and a sufficient oxygen concentration, above the hypoxia limit, leading to four possible conditions at any point in the tissue: 1) *C*_T-DM1_ *>* IC_50_ and 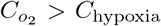 2) *C*_T-DM1_ *>* IC_50_ and 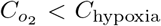, 3) *C*_T-DM1_ *<* IC_50_ and 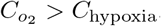, and 4) *C*_T-DM1_ *<* IC_50_ and 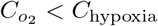

The Student’s t-test is used to test if the difference in mean of two distributions is statistically significant. The null hypothesis of the test is that the mean of the two distributions is the same. If the null hypothesis is rejected, that is the p-value from the test is below 0.05, the two distributions have a different mean.

Two methods are used to quantify how well the drug and oxygen concentrations overlap. Firstly, the overlap index is used [39],

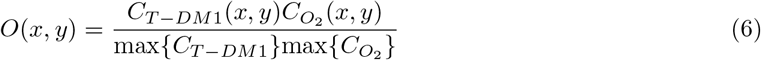

where *O*(*x, y*) is the overlap index at position (*x, y*), *C*_*T −DM*1_(*x, y*) and 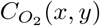 are the T-DM1 and oxygen concentrations at position (*x, y*), and max {*C*_*T −DM*1_} and max 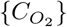 are the maximum T-DM1 and oxygen concentrations in the tissue. The overlap index is bound between 0 and 1, where 0 means at least one of the two is absent, whereas 1 indicates both are present in their maximum quantities. Secondly, we use Moran’s Bivariate I to quantify the spatial correlation of oxygen with the drug, where Moran’s bivariate I is the correlation coefficient of a variable in space with the weighted average of the other variable in the neighbouring positions [40]. A Moran’s I of 0 indicates the two variables are uncorrelated, 1 that they are perfectly positively correlated, and -1 that they are perfectly negatively correlated [40].

## Results

### Compression reduces tissue oxygenation

We start by investigating the effect that the tumour model (vessel compression) has on the fraction of hypoxic tissue. Figure 2a–b shows heat maps for the oxygen concentration in the control model and tumour model, respectively. The heat maps reveal that, in the control model, most of the low oxygen concentration areas are due to avascular areas in the network. On the contrary, in the tumour model, one can see that the core region also has low oxygen concentration levels. In effect, in the core region the mean oxygen concentration falls from 10.85 mmHg (median 11.17 mmHg) to 6.24 mmHg (median 6.04 mmHg) in the tumour model, while the tumour model has a more heterogeneous oxygenation with an interquartile range of [3.62 mmHg, 8.33 mmHg] compared to the control model interquartile range of [9.77 mmHg, 12.30 mmHg]. In addition to the increased heterogeneity, the oxygen concentration interquartile range in the tumour model is entirely hypoxic (*<* 8 mmHg), contrary to the control model. The distribution of oxygen concentrations in the tumour model and control model are plotted in Figure 2d, showing a lower mean and wider distribution in the tumour model compared to the control model (difference is statistically significant).

**Figure 2.**
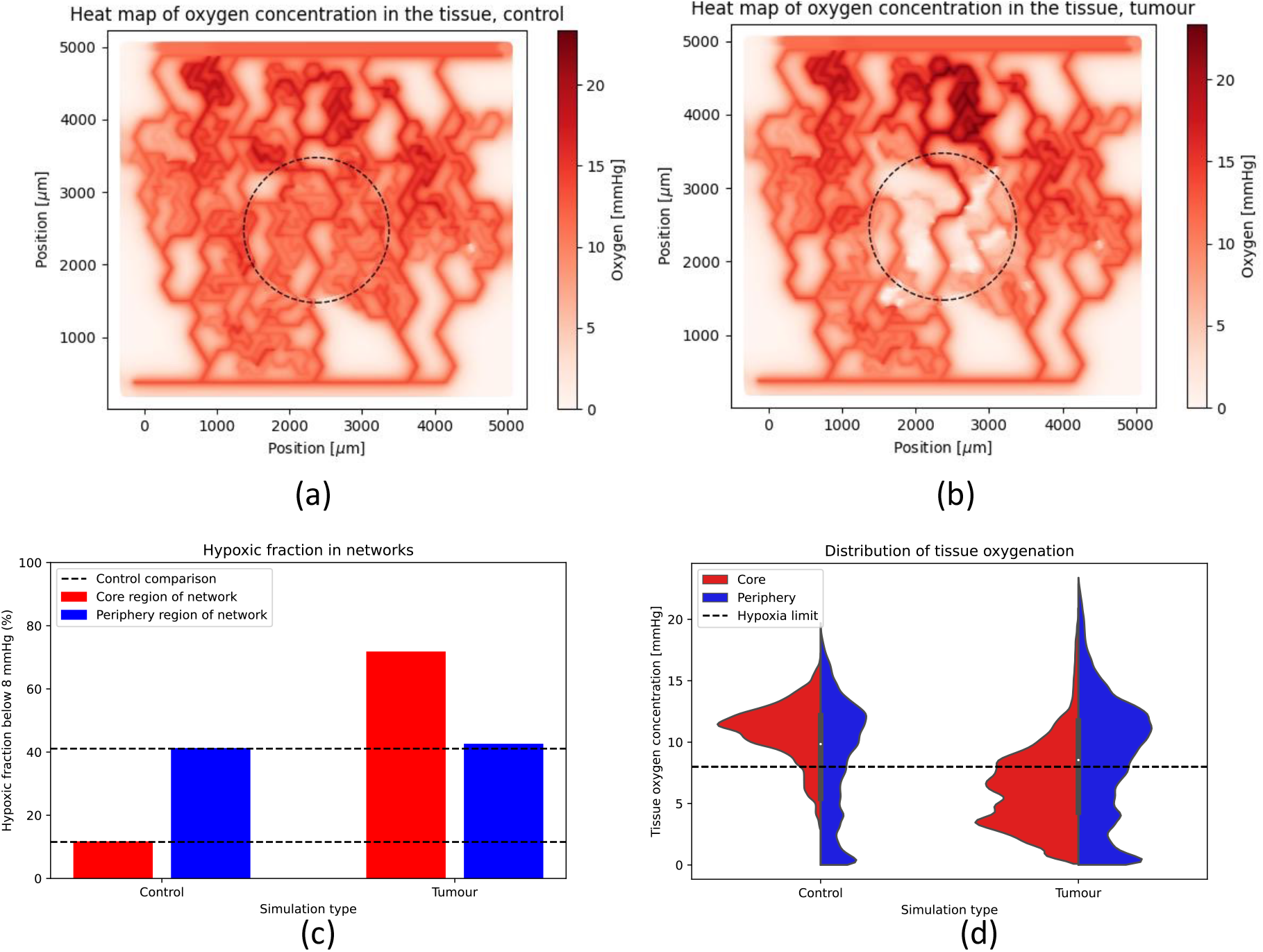
(a) and (b) show tissue oxygenation in mmHg for the control model (a) and the tumour model (b), the dashed circle delineates the core and periphery regions. (c) shows the fraction of hypoxic tissue in each model, in the core and periphery regions. (d) shows violin plots for each model, in the core and periphery regions, dashed line is the hypoxia limit at 8mmHg.

We can observe from Figure 2a–b that the periphery region of the tumour model also contains changes in the oxygenation compared to the control, particularly downstream of the compression. However, the mean oxygenation in the periphery of the control, 8.51 mmHg, is similar to that of the tumour model, 8.57 mmHg. The difference between the two distributions is statistically different, but the difference between the two means is 0.01 of a standard deviation, signifying the effect size is almost inexistent.

We next interrogate what fraction of the core region is hypoxic, defined as having an oxygen concentration below 8 mmHg of oxygen. In the control, the hypoxic fraction in the corw area is 11.5%, rising to 71.6% in the tumour model, representing a majority of the tissue. While in the periphery, the hypoxic fraction is similar in the control and in the tumour models, with the hypoxia being attributed to the aforementioned avascular areas. We hypothesise that the difference in hypoxic fraction between the control and the tumour models in the core region will reduce the efficacy of T-DM1, as T-DM1 is not as efficacious in hypoxic environments.

### Drug and oxygen overlap poorly and hypoxia reduces drug efficacy

Next, we investigate how the changed oxygenation of the tissue due to the tumour (compressed vessels) has an effect on the efficacy of T-DM1. Figure 3a–b shows binary heat maps indicating where there is both sufficient oxygen and sufficient T-DM1 to have a good drug effect. The heat maps clearly show a reduced amount of tissue fraction having enough of both oxygen and T-DM1 in the tumour model, falling from 82% to 28% of the tissue, Figure 3c–d. We further investigate if that change in T-DM1 efficacy is changed by the IC_50_ value used, and the results, supplementary Figure S5, shows that the results with an IC_50_ of 6.8 nM are similar to that of an IC_50_ of 2.9 nM, Figure 3.

**Figure 3.**
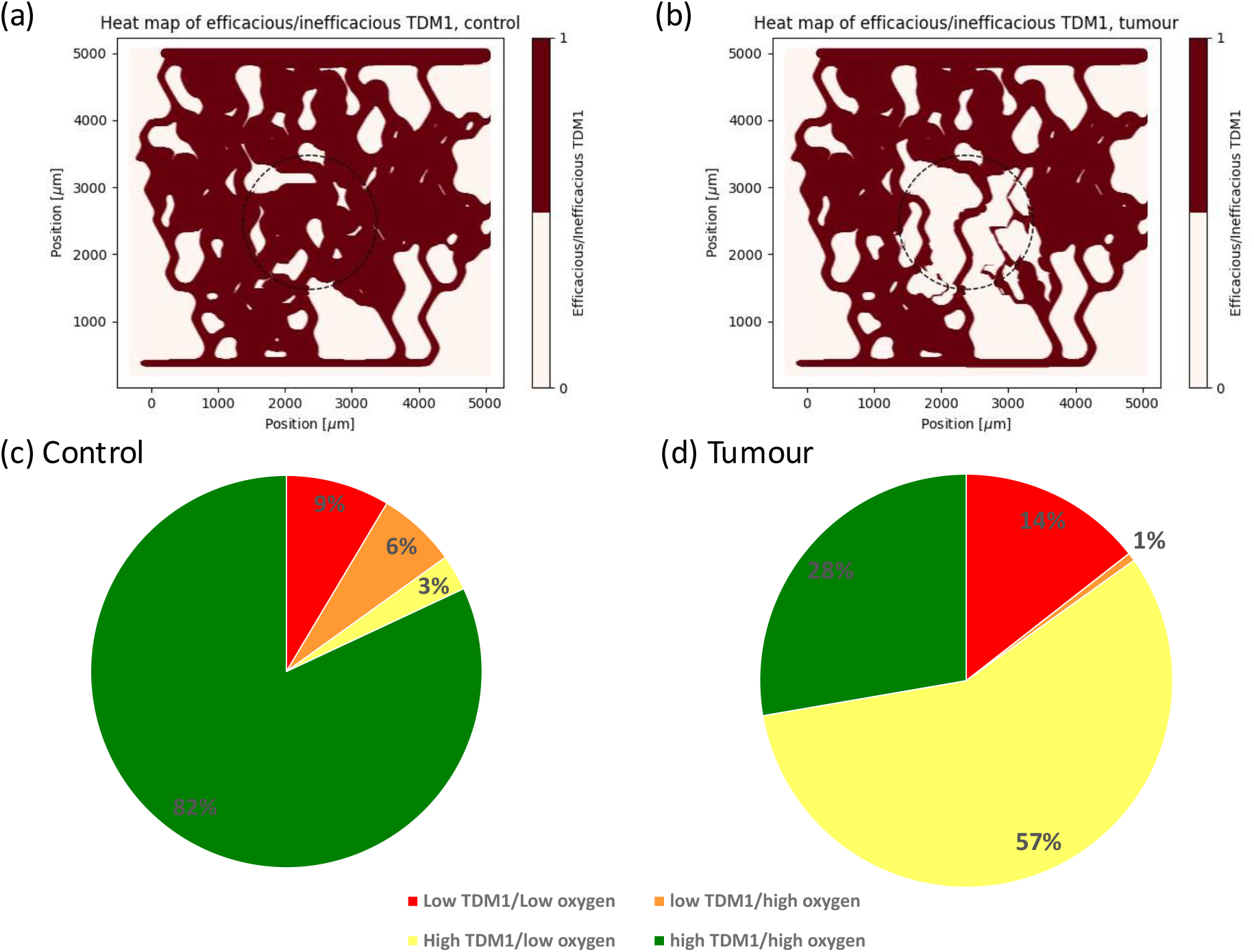
(a) Shows efficacious T-DM1 (oxygen above 8mmHg and T-DM1 above 2.9 nM, both required for internalisation and killing cells, respectively) in control simulation. (b) Shows efficacious T-DM1 in tumour simulation. (c) Shows fraction of tissue in core region corresponding to sufficient/insufficient oxygen/T-DM1 in control (c) and tumour (d) cases.

We then investigate what is the cause of the poor effect of T-DM1 in the tissue. Figure 3c–d breaks down the effect of the drug in the tissue into its four components (with respect to sufficient oxygen and sufficient T-DM1, see methods for more details). It reveals that the loss in T-DM1 efficacy is mostly due to the insufficient oxygen for the T-DM1 to internalise properly, with 3% of the tissue in the control case having sufficient T-DM1 but insufficient oxygen, compared to 57% in the tumour case.

Finally, we use two metrics to calculate the spatial correlation of oxygen concentration and T-DM1 concentration, Moran’s bivariate I and the overlap index, see methods for more details. In the control case, Moran’s bivariate I is 0.59, falling to 0.27. While the overlap index falls from a mean of 0.18±0.21 in the control case to 0.09±0.12 in the tumour model, the difference is statistically significant. The reduction in both Moran’s I and the overlap index illustrates that the tumour model, through compressing vessels, leads to a lower spatial correlation of oxygen concentration and drug concentration.

### The effect of inlet haematocrit and T-DM1 depletion from blood stream

One might speculate that the extent of hypoxia can simply be alleviated via increasing the inlet discharge haematocrit which could in turn improve the efficacy of T-DM1. In order to assess its impact on T-DM1 efficacy, in Figure 4a–b we vary the inlet discharge haematocrit between 5 and 30%. Inlet discharge haematocrit below 15% is too low to produce sufficient oxygenation even in the close promixity of the blood vessels, independent of the compression status, which results in almost the entire tissue being hypoxic (Figure 4a). Above this threshold, increasing inlet discharge haematocrit indeed yields an improved drug efficacy. However, while in the control case the efficacy improves greatly so that for 25-30% inlet discharge haematocrit, the majority of the tissue experiences sufficient concentrations of oxygen and T-DM1, the tumour model results in the majority of the tissue remaining inefficaciously perfused even for 30% inlet discharge haematocrit. The spatial correlation between oxygen and T-DM1, as evaluated by Moran’s I, only varies mildly as inlet discharge haematocrit is increased independent of tumour status (Figure 4b). Therefore, the weak correlation observed when vessels are compressed cannot simply be rectified by increasing the inlet discharge haematocrit.

**Figure 4.**
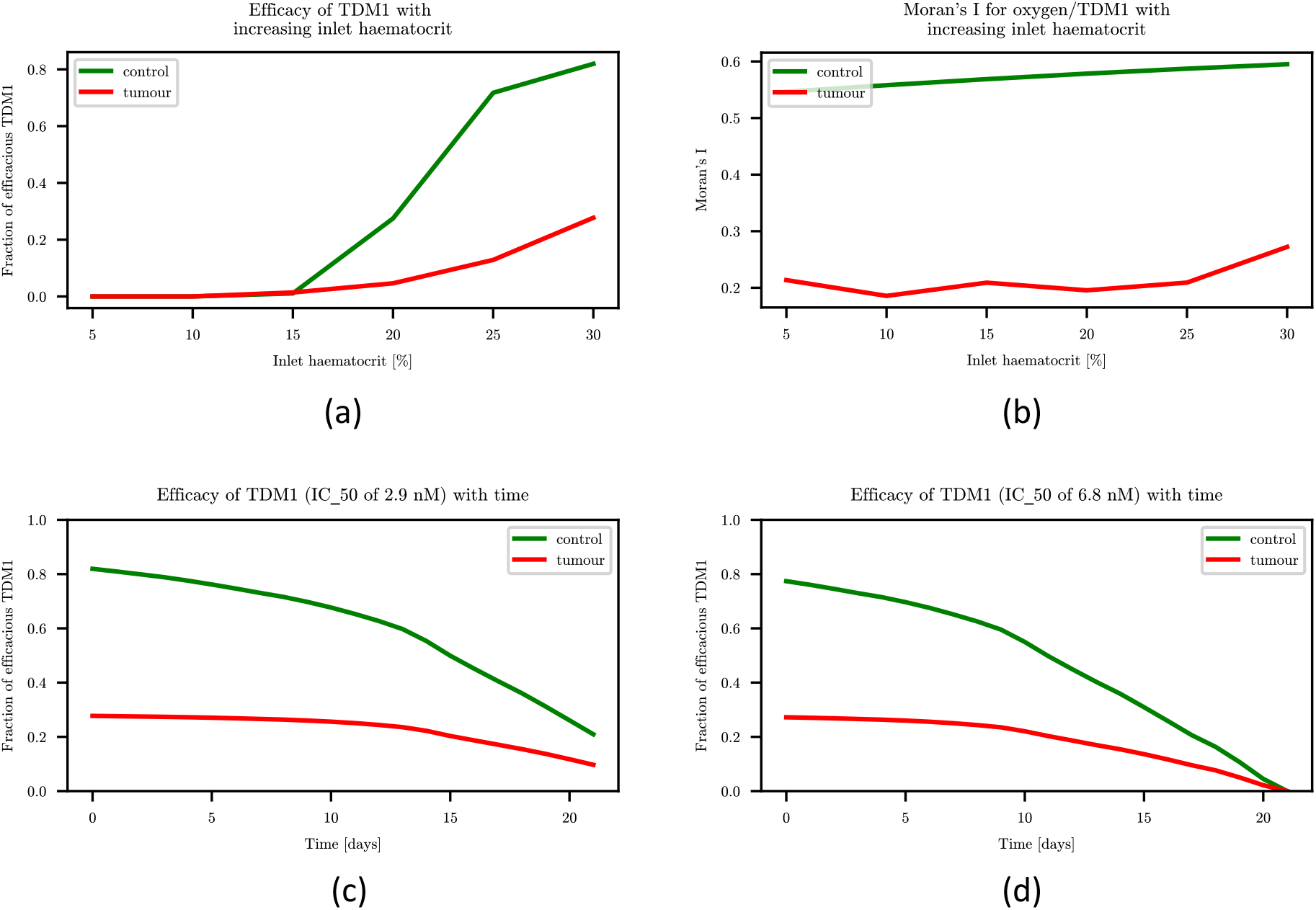
(a) shows how increasing inlet discharge haematcorit changes the fraction of high oxygen/high T-DM1 in the tissue. (b) shows how Moran’s I changes with inlet discharge haematocrit. (c) shows efficacy of T-DM1 with time when IC_50_ = 2.9 nM. (d) shows efficacy of T-DM1 with time when IC_50_ = 6.8 nM.

Motivated by the rate at which T-DM1 concentration in the blood stream decays, drug injections are standardly administered every 3 weeks [37] (see also Figure S1). Figure 4c–d documents that due to the decay of T-DM1 from plasma, the efficacy of drug in the tissue decreases with time to a point where only a negligible fraction of the tissue experiences efficacious T-DM1, using two thresholding values – IC_50_ = 2.9 nM (panel c) and IC_50_ = 6.8 nM (panel d) – for two cell lines studied in [10]. Interestingly, in the tumour model when vessels are compressed, the fraction of tissue experiencing efficacious T-DM1 is low and essentially independent of the particular value of IC_50_. Confirming again that the primary factor preventing efficacious treatment is the prevalence of hypoxia caused by the tumour induced vessel compression.

## Discussion

The tumour microenvironment acts as a barrier to drug delivery and leads to tumour tissue hypoxia. In addition, some drugs require the presence of oxygen to be fully functional. We therefore hypothesised that the TME leads to areas of high drug concentration poorly overlapping with areas of high oxygen concentration due to their different modes of transport to tissue. In this work we tested this hypothesis with a computational model of a tumour with compressed tumour blood vessels, with oxygen and T-DM1 (an oxygen-dependent drug) transport from the blood vessels to the tissue, compared to a control model without compressed vessels.

We started by deriving a novel model reduction for T-DM1 transport based on key timescales, simplifying its numerical solution (it is worth noting that the model reduction derived in this work for T-DM1 might not be applicable for other drugs; this will always depend on the relevant timescales of transport and uptake processes [5]). We then simulated blood flow through tumour induced compression, and control, vessels which then acted as sources of both oxygen and T-DM1 to diffuse to the tissue. As the tumour compressed vessels introduce haematocrit heterogeneity in the microvascular networks, the tumour tissue oxygenation was found to be more heterogeneous than the control, with a hypoxic fraction increasing from 11.5% to 71.6%. We then showed that due to the higher fraction of hypoxia in the tumour tissue model, there is a lower fraction of efficacious T-DM1, 28% compared to 82% in a control model, as its internalisation is impeded in hypoxic conditions. This reduction in efficacious T-DM1 is the result of a reduced overlap of areas of high oxygen concentration with areas of high T-DM1 concentration. Finally, we showed that increasing the inlet discharge haematocrit to the network is insufficient to improve the efficacy of T-DM1 to levels comparable to that of a control.

One of the implications of this work is that independently considering oxygen transport and drug transport in oxygen dependent drugs can lead to lower drug efficacy than expected. Given that multiple important drugs, such as doxorubicin [11] and T-DM1 [10] amongst others [13], require the presence of oxygen to function, this work suggests that the overlap of drug and oxygen is an important biomarker for drug efficacy and therefore treatment outcome. We use tumour-induced vessel compression [27, 41] and T-DM1 [37] as exemplars of a vascular abnormality and drug transport, and hypothesise that this effect would occur with other structural abnormalities; such as decreased inter-bifurcation length [17, 20], increased tortuosity [42] and leakiness [43]; and other drugs with different transport properties [5].

Future work combining experimental validation and numerical modelling could enable one to predict what type of abnormalities present in a tumour microvascular network would lead to poor oxygen/drug overlap depending on the transport properties of a specific drug. One could then characterize the transport properties of microvascular networks via constructing governing dimensionless groupings which incorporate all key model parameters such as the vascular density, the inlet discharge haematocrit as well as other parameters determining the oxygen and drug perfusion (see [36] for the former and Table S1 for the latter). We expect that this approach would provide more robust insights into drug efficacy assessment (e.g. a phase diagram dividing the parameter space into subregions where drug or oxygen perfusion limits the treatment efficacy) independent of a particular network design.

Finally, normalisation therapy offers a means of therapeutically changing tumour microvascular networks, with the aim of temporarily making them more efficient [44, 45]. For example, angiotensin inhibition decompresses vessels and improves tissue oxygenation [44], through the opening of blocked vessels [44] and haematocrit homogenisation [18]. However, despite its promising approach, normalisation therapy has remained challenging [46]. This work postulates that normalisation could aim to improve drug and oxygen overlap, which could be tested experimentally, to improve therapeutic efficiency.

## Acknowledgments

M.O.B. gratefully acknowledges funding from: Fondation Leducq Transatlantic Network of Excellence (17 CVD 03); EPSRC grant no. EP/X025705/1; British Heart Foundation and The Alan Turing Institute Cardiovascular Data Science Award (C-10180357); Diabetes UK (20/0006221); Fight for Sight (5137/5138); the SCONe projects funded by Chief Scientist Office, Edinburgh & Lothians Health Foundation, Sight Scotland, the Royal College of Surgeons of Edinburgh, the RS Macdonald Charitable Trust, and Fight For Sight; the Neurii initiative which is a partnership among Eisai Co., Ltd, Gates Ventures, LifeArc and HDR UK. J.B. was funded by the Medical Research Council Precision Medicine Doctoral Training Programme scholarships (MR/N013166/1).

## Supplementary material

### Blood flow model

#### Physical model

Blood is treated at each vessel segment as a one-dimensional continuous and incompressible fluid with an apparent viscosity, this is done through the use of Poiseuille’s law [15]:

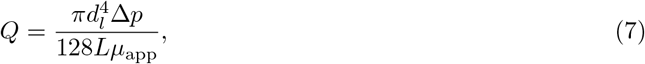

where *Q* is the flowrate of blood at each vessel segment, *d*_*l*_ is the vessel diameter of the vessel segment, Δ*p* is the pressure drop along the vessel segment, *L* is the length of the vessel segment, and *µ*_*app*_ is the apparent viscosity of blood in the vessel segment. Poiseuille’s law can also be expressed for an elliptical cross-section when vessels are compressed [18]:

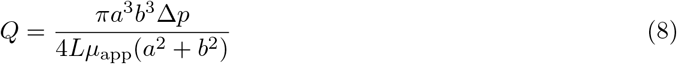

where *a* and *b* are the major and minor radius of the ellipse, respectively [18]. If *a* = *b*, the elliptical formulation of Poiseuille’s law, Eq. 8 is the same as the circular one, Eq. 7.

The apparent viscosity of blood in each vessel segment is calculated using an empirically derived relationship where the apparent viscosity is dependent on vessel diameter and discharge haematocrit [31, 32]:

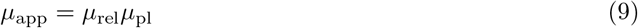

where *µ*_*pl*_ is the apparent viscosity of pure plasma and *µ*_*rel*_ is the relative apparent viscosity, defined through

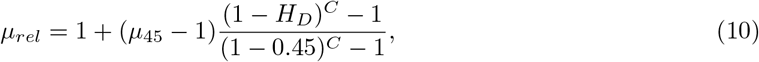

where *µ*_45_ is the apparent viscosity of blood at a discharge haematocrit, *H*_*D*_, of 45%, itself defined through the following relation

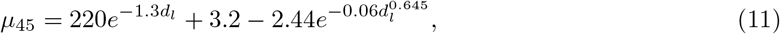

where *d*_*l*_ is the diameter of the vessel segment in microns. *C* is defined through

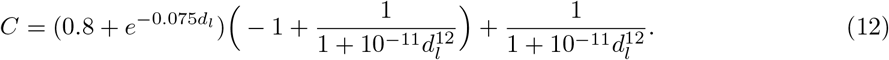

This system of equations can be solved with known boundary conditions (either pressure or velocity boundary conditions) and if the discharge haematocrit is known at every vessel segment [47]. However, the discharge haematocrit is not known a priori, as red blood cells are distributed in networks disproportionately to blood flow [15]. The following relation allows one to predict the partitioning of red blood cells at each diverging microvascular bifurcation once the flowrates in the network are known

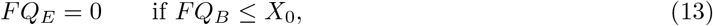

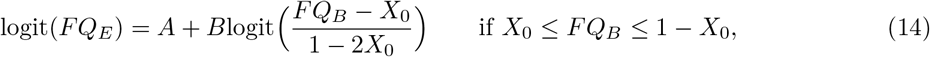

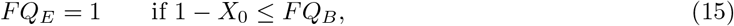

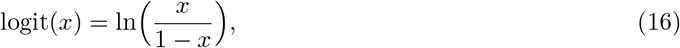

where *FQ*_*E*_ is the flowrate fraction of red blood cells from the parent branch that flows to the daughter branch, and *FQ*_*B*_ is the fractional flowrate of blood from the parent branch that flows to the daughter branch. *A, B*, and *X*_0_ are further defined by the following relationships

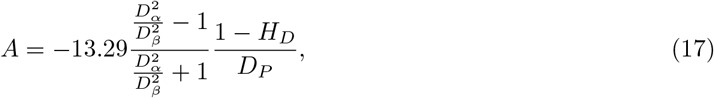

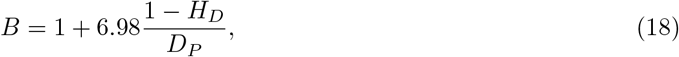

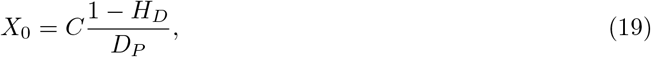

where *D*_*α*_ is the diameter of the child branch for which *FQ*_*E*_ and *FQ*_*B*_ are calculated, *D*_*β*_ is the diameter of the other child branch, *D*_*P*_ is the parent branch diameter, and *H*_*D*_ is the parent branch discharge haematocrit. The value of *C* depends on whether the parent branch is compressed and *C* = 0.96 for circular (non-compressed) vessels [32], while *C* = 4.16 for compressed vessels with an aspect ratio of 4.26 [18], as assumed in this work. This leaves only the discharge haematocrit of inlets to be defined as a final boundary condition to solve the system of equations.

#### Numerical model

The physical model described above needs to be numerically solved. The solution is obtained through a previously described iterative scheme [47, 48], necessary due to the flow dependent disproportional partitioning of red blood cells in Eqs. 13–15. At each iteration of the solver, the system of Poiseuille equations, Eqs. 7–8, is solved with the known boundary conditions. With the known flowrate in each vessel, the discharge haematocrit is calculated in each vessel, Eqs. 13–15, where the discharge haematocrit in the inlet vessels is also a fixed boundary condition. The solution for the flowrate of both blood and red blood cell flow at each iteration is compared to the solution at the previous iteration is then compared.

The solver is considered to have converged when

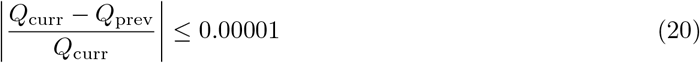

where *Q*_*curr*_ and *Q*_*prev*_ are the values in the current and previous iteration of a given flowrate (blood or red blood cell) in a vessel. The iterative process continues until all vessels have converged.

The solver is implemented in a custom python2.7 code.

**Figure S1.**
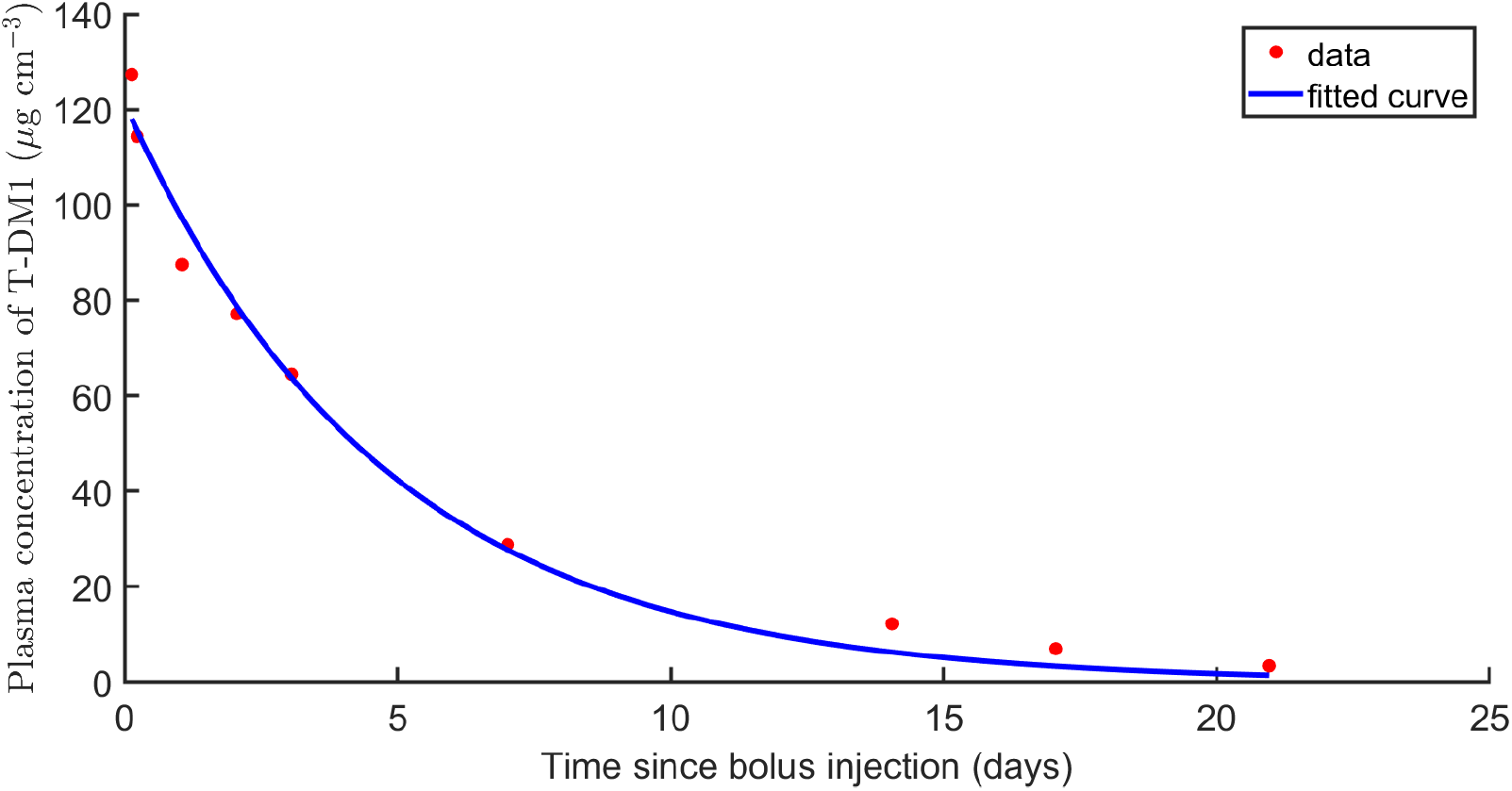
Time evolution of plasma concentration using the highest-dose from [37] (reproduced from Figure 1a therein) and the corresponding single-term exponential fit.

**Table S1.**
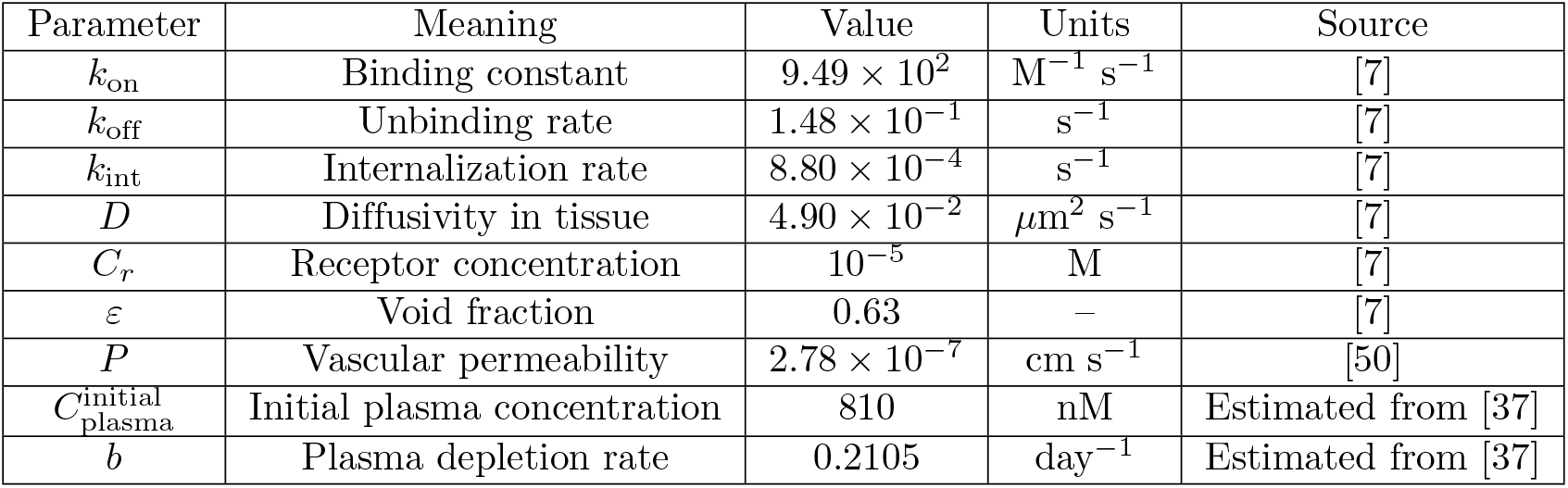
Parameters for T-DM1 simulations.

### Reduced-order model for drug perfusion

#### Estimation of dimensional parameters

The majority of parameters governing pharmacokinetics of T-DM1 are readily available from [7]: see Table 1 (non-FUS fitted parameters) for the void fraction (referred to as “interstitium effective porosity”) and Table 2 (non-FUS fitted parameters) for the other parameters. As a trastuzumab molecule is much larger than its emtansine counterpart (as measured by the respective molecular weights: 738 Da for DM1 and 145,167 Da for trastuzumab [49]), we estimate the vessel permeability for T-DM1 with the value for trastuzumab, i.e. *P* = 10^*−*3^ cm h^*−*1^ = 2.78 *×* 10^*−*7^ cm s^*−*1^ [50]. To fully specify the model, we need to provide a relationship describing the depletion of T-DM1 from the blood stream, i.e. *C*_plasma_(*t*). To this end, we reproduce the highest-dose (4.8 mg/kg) data from Figure 1a in [37], using linear scaling for the *Y* axis and find the best single-term exponential fit to the data using MATLAB’s fit() functionality. The best fit is shown in Figure S1. The resulting single-term exponential fit is summarized in Eq. (3). Note that we used the molecular weight of T-DM1 from [49] (*≈* 1.5 *×* 10^5^ Da, as discussed above) to convert the units (*µ*g to mol); this way the fitted initial (for instance) plasma concentration of 121.5 *µ*g cm^*−*3^ is converted to 8.1 *×* 10^*−*10^mol cm^*−*3^ = 810 nM. All dimensional models parameters are summarized in Table S1.

#### Nondimensionalization

We nondimensionalize concentrations with respect to the (constant) receptor concentration *C*_*r*_, lengths with respect to a typical, for a randomly selected location in the tissue domain, distance to the nearest vessel *s*_min_ (here estimated as 50 *µ*m) and time with respect to the plasma depletion timescale (the only slow timescale independent of locally-varying quantities – vessel diameter and inter-vessel distance), i.e.

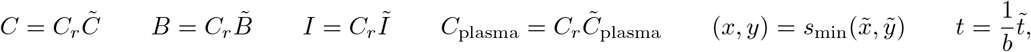

where the dimensionless quantities are denoted with tildes. Recalling that the units of Dirac delta function equals one over the units of its argument, we get from Eqs. (2) after algebraic manipulation the following dimensionless model

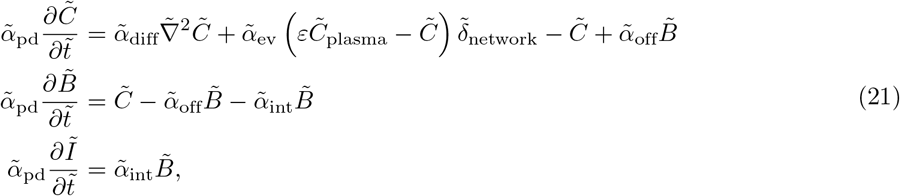

where

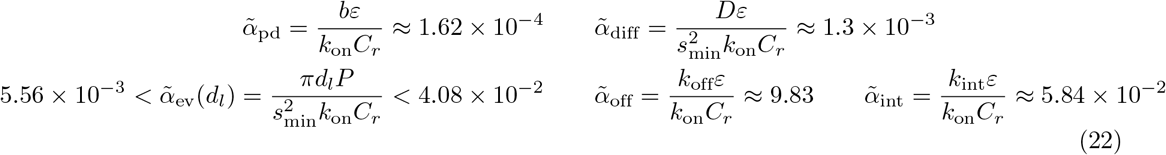

denote dimensionless parameter groupings indicating plasma-depletion, diffusion, extravasation, unbinding and internalization rates relative to the binding rate *k*_on_*C*_*r*_*/ε*. Note that these dimensionless groupings can alternatively be expressed in terms of standard dimensionless numbers governing pharmacokinetics, such as Biot and Damkohler numbers [5]. Finally, the dimensionless counterpart of the plasma depletion Eq. (3) reads

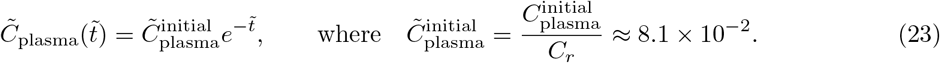

#### Model reduction

By considering the estimates in Eq. (22), the time derivative in the second equation in Eq. (21) can be neglected, yielding the first quasi-steady approximation

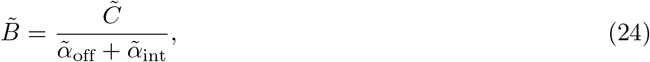

which upon redimensionalizing gives Eq. (4). Substituting Eq. (24) back into the first equation in Eq. (21) gives

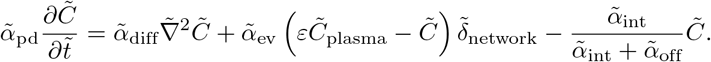

From the estimated values in Eq. (22), we get 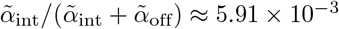 and we conclude that the time-derivative term can be neglected yielding the final reduced-order model

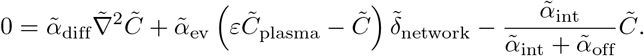

Upon redimensionalizing, we arrive at Eq. (5) which is solved in Microvessel Chaste [34].

### Simulations on simple geometry confirm the validity of the reduced-order model

#### Simple geometry: model and boundary conditions

To confirm the validity of above-derived approximation, we propose the same problem on a simple geometry containing only a single, infinitely-long, cylindrical blood vessel of diameter *d*_*l*_ with the axis oriented in the *Z* direction. This vessel supplies the drug to a surrounding tissue domain in the form of a cylindrical annulus of an inner diameter *d*_*l*_ and an outer diameter equal to an inter-vessel distance *h*_*l*_, where both *d*_*l*_ and *h*_*l*_ cover ranges found in our networks. Provided no variation in drug concentration in the *Z* direction (i.e. along the vessel) is introduced via initial or boundary conditions, we can assume that the resulting drug profile remains axisymmetric, i.e. *C*(*t, x, y, z*) = *C*(*t, r*), where 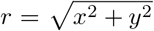 denotes the polar radius. Importantly, the Dirac delta source term in Eq. (2) is here replaced by the boundary condition

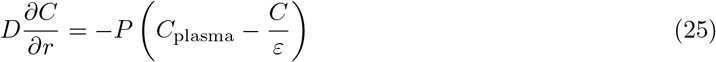

imposed at *r* = *d*_*l*_*/*2 and we also impose zero-flux boundary condition

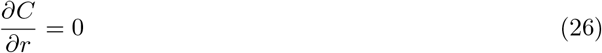

at *r* = *h*_*l*_*/*2. The full model thus reads

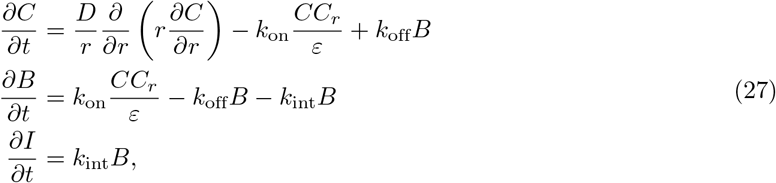

whereas the reduced-order model reads

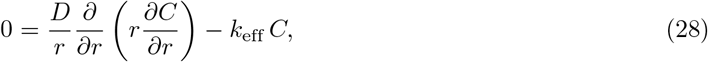

where *k*_eff_ is defined in Eq. (5), supplemented by Eq. (4) and the third equation in Eq. (2) governing *B* and *I*, respectively. Both problems are to be solved on a spatial domain *d*_*l*_*/*2 ≤ *r* ≤ *h*_*l*_*/*2 subject to boundary conditions, Eq. (25) and Eq. (26), and the initial conditions

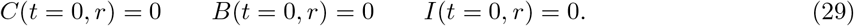

#### Analytic solution of the reduced-order problem

Due to its linearity, the reduced-order problem can (when posed on the simple geometry) be solved directly. Multiplying Eq. (28) by *r*^2^ and dividing by *D* we get

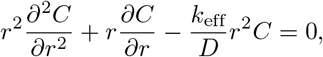

Denoting 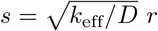 *r* and *F* (*t, s*) = *C*(*t, r*), this problem transforms to

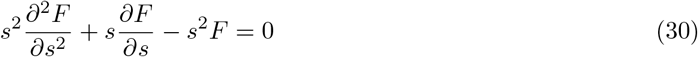

which is to be solved in the domain 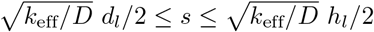 subject to boundary conditions

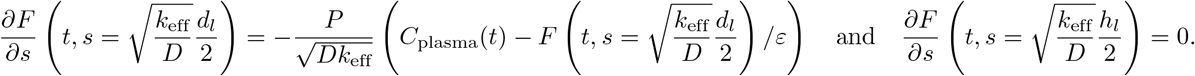

For any fixed time *t*, Eq. (30) is a modified Bessel equation of order 0 and the solution is therefore a linear combination of its two linearly independent solutions *I*_0_(*s*) and *K*_0_(*s*) (modified Bessel functions of the first and second kind), i.e.

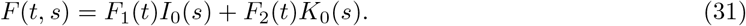

Letting ‘ denote the derivative with respect to *s*, the two boundary conditions then give (for arbitrary fixed time *t*) a system of two linear equations for the unknown *F*_1_(*t*) and *F*_2_(*t*) of the form

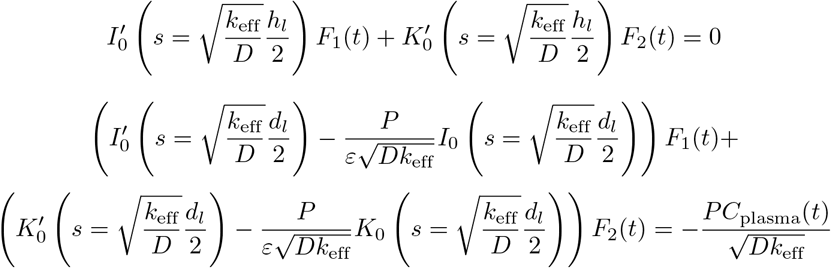

Finally, using known properties of modified Bessel functions, we have

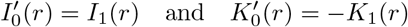

for any *r*. This system can easily be solved and *F*_1_(*t*) and *F*_2_(*t*) obtained for arbitrary *t >* 0.

#### Numerical method for the solution of the full model

Recall the full dimensional problem Eq. (27) which is to be solved on a one-dimensional spatial domain *d*_*l*_*/*2 ≤ *r* ≤ *h*_*l*_*/*2 over time interval 0 ≤ *t* ≤ *T* (where *T* = 21 days), subject to initial conditions Eq. (29) and boundary conditions Eqs. (25)-(26). We solve this problem numerically using finite difference schemes, using standard centered finite differences for spatial derivatives (including in the boundary conditions) and the (implicit) backwards Euler method for temporal derivatives. We subdivide the spatial domain into *N* intervals of equal length Δ*r* = (*h*_*l*_ *− d*_*l*_)*/*(2*N*) by the set of *N* + 1 spatial points

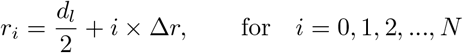

so that *r*_0_ = *d*_*l*_*/*2 and *r*_*N*_ = *h*_*l*_*/*2. Similarly, we subdivide the temporal domain into *M* intervals of equal length Δ*t* = *T/M* by the set of *M* + 1 temporal points

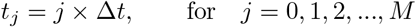

so that *t*_*M*_ = *T* . We then denote the numerical approximations at selected temporal (*j*) and spatial (*i*) points as

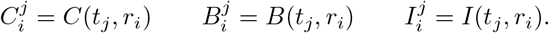

#### Discretization of the differential equations

Below, we will use Δ*r*^2^ to denote (Δ*r*)^2^. Centering at *r* = *r*_*i*_, we apply the finite differences to the differential equations, which gives

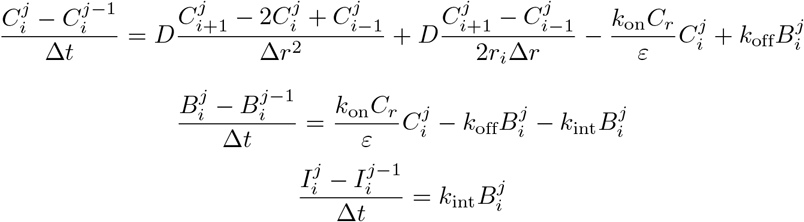

which upon rearrangement leads to

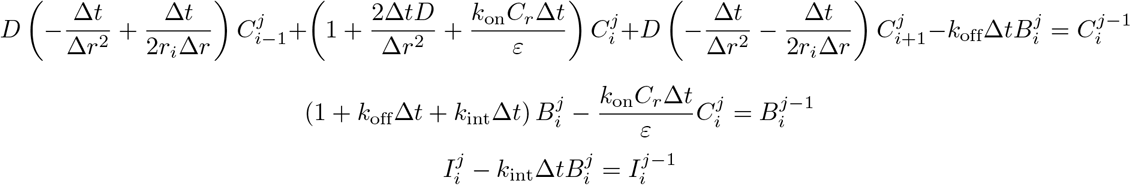

From the known initial conditions (*j* = 0), we can use these equations iteratively to calculate numerical solutions up to the final time *j* = *M* . These equations in their current form are valid for *i* = 1, …, *N −* 1. However, the equation for *C* centered at *d*_*l*_*/*2 requires the knowledge of non-existing *r*_*−*1_ and that centered at *r*_*N*_ requires the knowledge of non-existing *r*_*N*+1_. Therefore, these two equations must be modified using information coming from the boundary conditions.

#### Boundary conditions

At the boundary points, we introduce ghost points at which the values are estimated from the discretized boundary conditions. Introducing auxiliary spatial points outside the domain, the discretized boundary condition at *r* = *r*_*N*_ reads

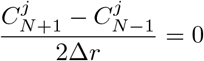

which gives

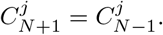

This can be substituted back into the corresponding (*i* = *N*) equation for *C* and we get

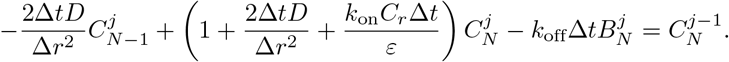

Similarly, the discretized boundary condition at *r* = *r*_0_ reads

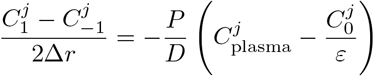

from which we conclude

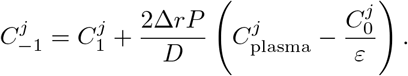

Substituting this back into the appropriate (*i* = 0) equation for *C*

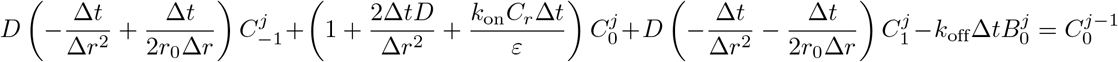

we get

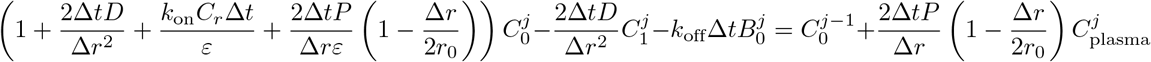

#### Formulating the linear system

*Ax* = *b* Thus, for every known solution (*C, B, I*) at time *j −* 1, we find the solution at time *j* by solving a linear system of algebraic equations of the form

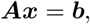

where we wish to find a vector

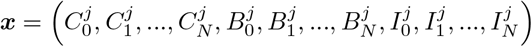

given a vector

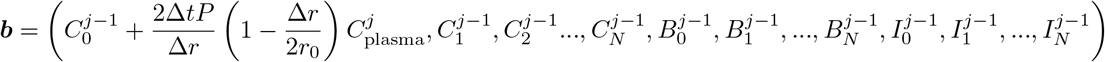

and ***A*** is a block matrix with 3 *×* 3 blocks of equal size, (*N* + 1) *×* (*N* + 1) (size 3(*N* + 1) *×* 3(*N* + 1) in total), which reads

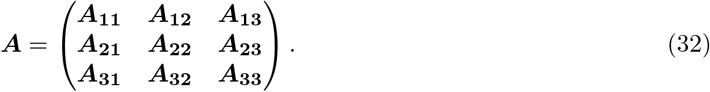

Detailing the blocks, we have

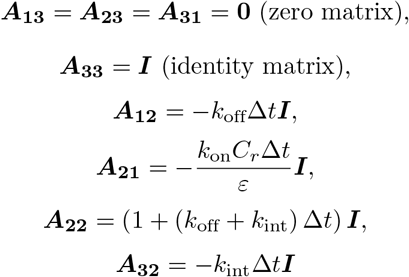

and

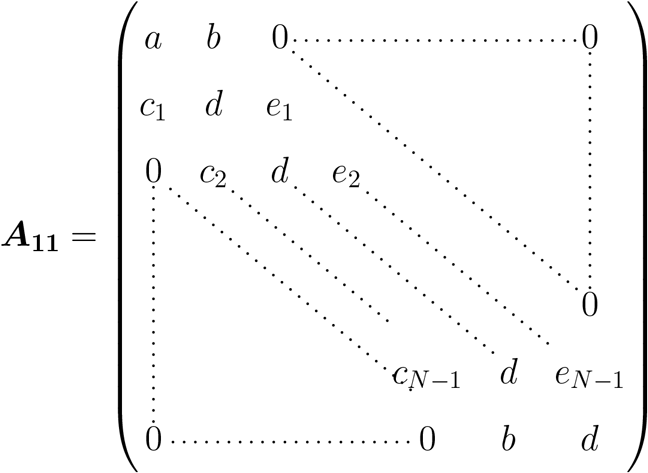

where

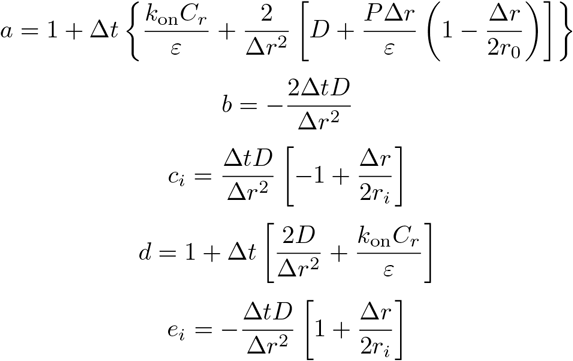

for *i* = 1, 2, …, *N −* 1. In MATLAB, we then solve *Ax* = *b* for unknown vector ***x*** using “x=linsolve(A,b)”.

**Figure S2.**
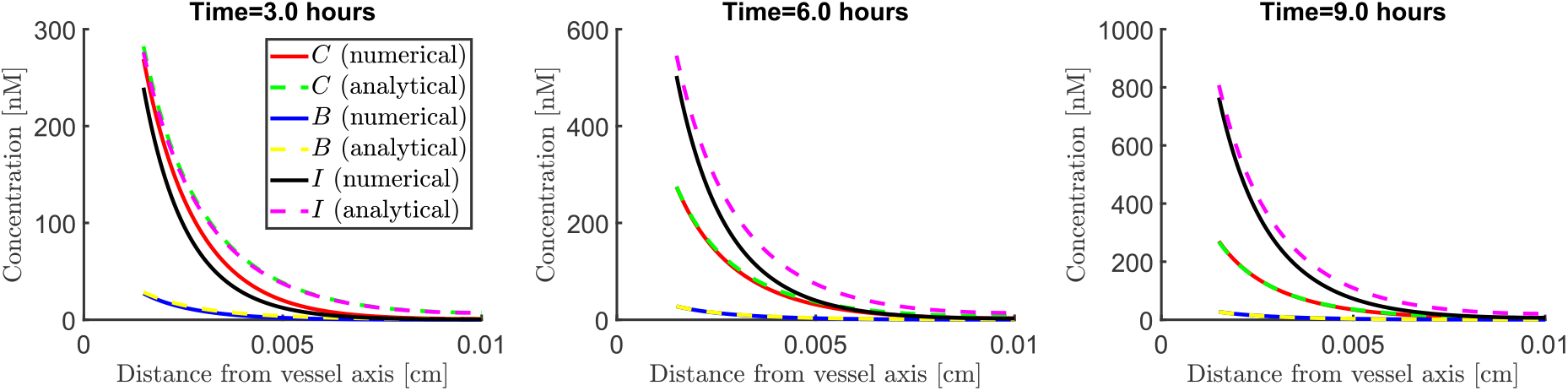
Comparison of solution profiles as functions of the radial coordinate *r* at early times *t* = 2, 4 and 6 hours after the bolus injection.

#### Validity of the new approximation

Finally, we need to check whether the analytical solution Eq. (31) provides a good approximation to the numerical solution of the full problem described in the previous section on the therapeutically relevant timescales (days to weeks). Using default parameter values from Table S1 and representative values of inter-vessel distance *h*_*l*_ = 200*µ*m and vessel diameter *d*_*l*_ = 30*µ*m, we find that early (3 hours) after the bolus injection, the reduced-order model gives a poor approximation to the full numerics (see Figure S2). However, after 6 *−* 9 hours, the reduced-order model already provides a very good approximation for free and bound drug concentrations. Due to the initial transient behaviour, which our reduced-order model does not capture, the prediction for the internalized drug *I* contains significant errors hours after the bolus injection. However, these differences become negligible as early as 1 day after the bolus injection (see Figure S3). Moreover, the reduced-order model performs remarkably well across the parameter space spanned by the inter-vessel distance *h*_*l*_ and the vessel diameter *d*_*l*_. As our analysis in the main body of this manuscript predicted, the largest errors occur for very large values of the vessel diameter and very small values of inter-vessel distance; however, 1 day after bolus injection, even these differences become small.

Estimation errors accumulated over the initial transient period become negligible especially over the timescale of weeks, as evidenced in Figure S4 (using representative model parameters). Note that as the plasma concentration decreases exponentially to 0, the amount of internalized drug increases at a declining rate. In summary, these figures confirm that the reduced-order model provides a useful approximation to the full-model behaviour on the timescales of interest (days to weeks).

**Figure S3.**
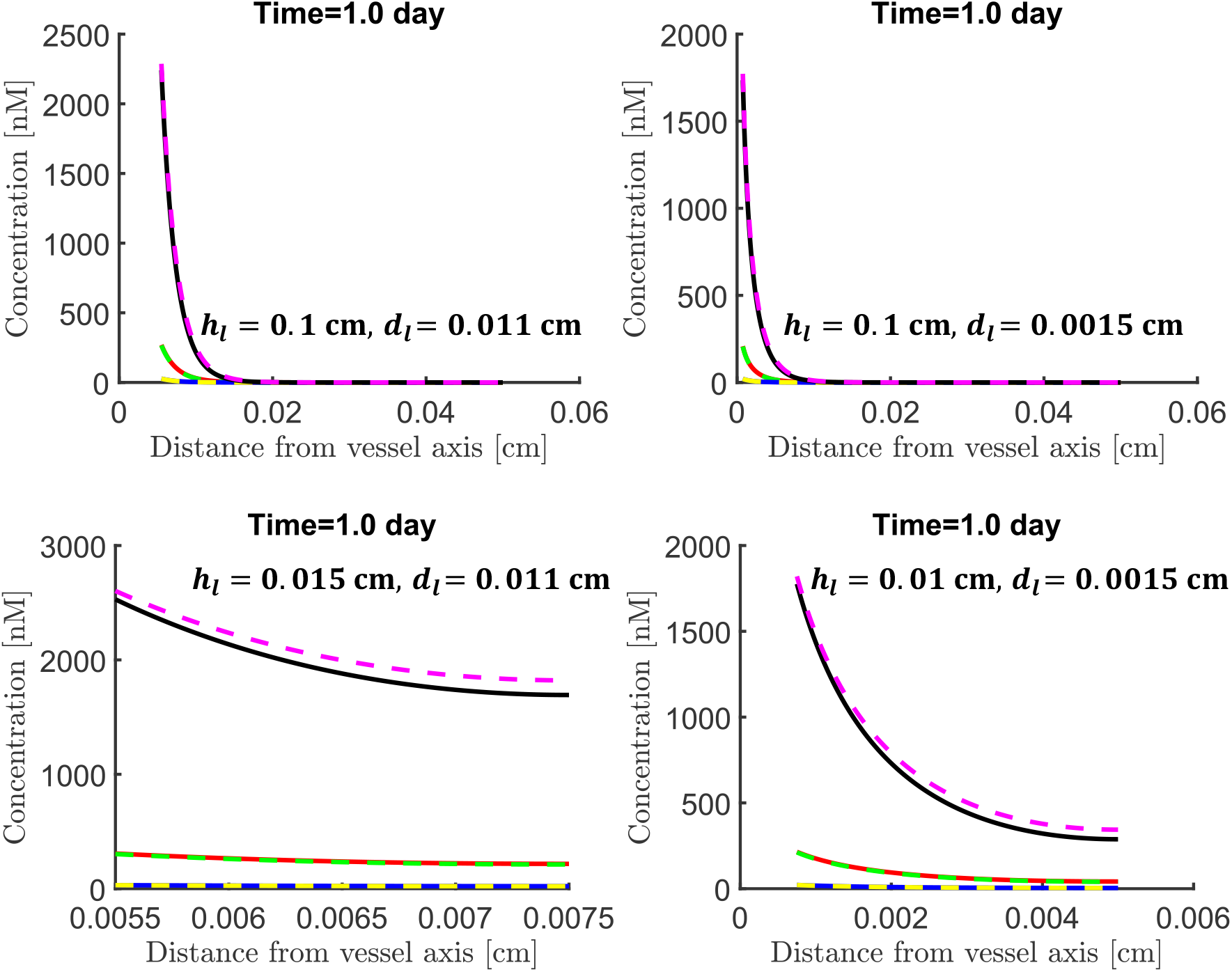
Comparison of solution profiles as functions of the radial coordinate *r*, 1 day after the bolus injection for different combinations of inter-vessel distance *h*_*l*_ and vessel diameter *d*_*l*_. The same legend as in Figure S2 applies.

**Figure S4.**
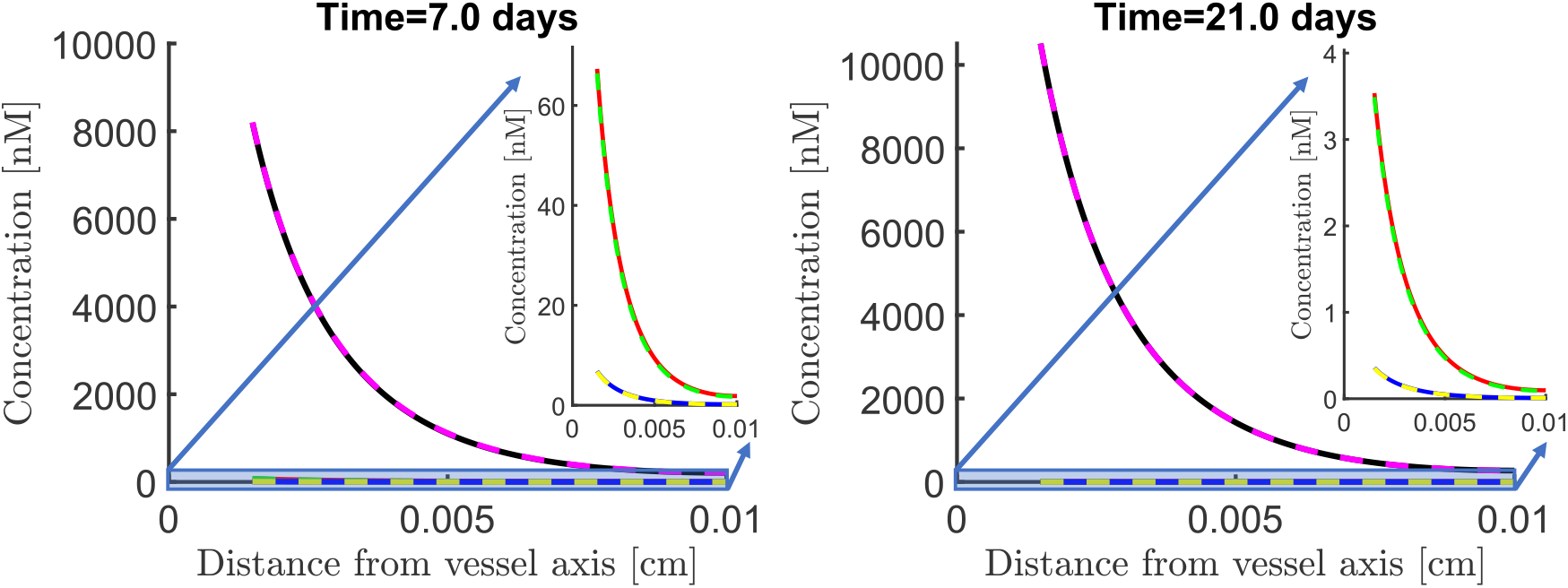
Comparison of solution profiles as functions of the radial coordinate *r* at times *t* = 7 and 21 days after the bolus injection. Insets on the right of each panel document the agreement for free drug concentration *C* and the bound drug *B*. The same legend as in Figure S2 applies.

**Figure S5.**
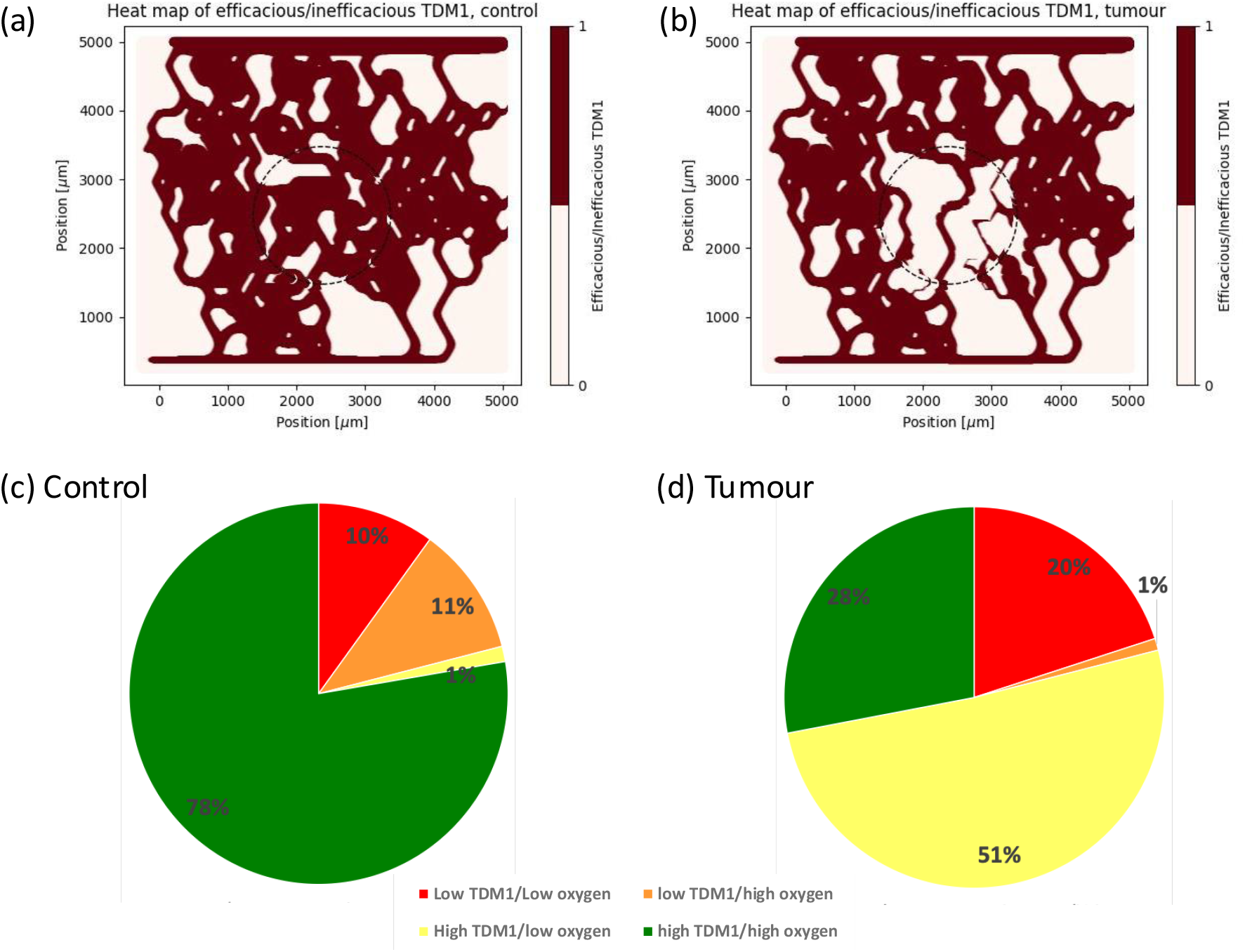
(a) Shows efficacious T-DM1 (oxygen above 8mmHg and T-DM1 above 6.8 nM, both required for internalisation and killing cells, respectively) in control simulation. (b) Shows efficacious T-DM1 in tumour simulation. (c) Shows fraction of tissue in core region corresponding to sufficient/insufficient oxygen/T-DM1 in control (c) and tumour (d) cases.

## References

1. Hanahan, D. & Weinberg, R. A. Hallmarks of cancer: The next generation Mar. 2011.

2. Dewhirst, M. W. & Secomb, T. W. Transport of drugs from blood vessels to tumour tissue. Nature Reviews Cancer 17, 738–750 (2017).

3. Stylianopoulos, T. & Jain, R. K. Combining two strategies to improve perfusion and drug delivery in solid tumors. Proceedings of the National Academy of Sciences 110, 18632–18637 (2013).

4. Dewhirst, M. W., Mowery, Y. M., Mitchell, J. B., Cherukuri, M. K. & Secomb, T. W. Rationale for hypoxia assessment and amelioration for precision therapy and immunotherapy studies. Journal of Clinical Investigation 129, 489–491 (Jan. 2019).

5. Thurber, G. M. & Weissleder, R. A systems approach for tumor pharmacokinetics. PLoS ONE 6 (2011).

6. Minchinton, A. I. & Tannock, I. F. Drug penetration in solid tumours Aug. 2006.

7. Arvanitis, C. D., Askoxylakis, V., Guo, Y., Datta, M., Kloepper, J., Ferraro, G. B., Bernabeu, M. O., Fukumura, D., McDannold, N. & Jain, R. K. Mechanisms of enhanced drug delivery in brain metastases with focused ultrasound-induced blood–tumor barrier disruption. Proceedings of the National Academy of Sciences 115, E8717–E8726 (2018).

8. d’Esposito, A., Sweeney, P. W., Ali, M., Saleh, M., Ramasawmy, R., Roberts, T. A., Agliardi, G., Desjardins, A., Lythgoe, M. F., Pedley, R. B., Shipley, R. & Walker-Samuel, S. Computational fluid dynamics with imaging of cleared tissue and of in vivo perfusion predicts drug uptake and treatment responses in tumours. Nature Biomedical Engineering 2, 773–787 (2018).

9. Tong, R. T., Boucher, Y., Kozin, S. V., Winkler, F., Hicklin, D. J. & Jain, R. K. Vascular Normalization by Vascular Endothelial Growth Factor Receptor 2 Blockade Induces a Pressure Gradient Across the Vasculature and Improves Drug Penetration in Tumors. Cancer Research 64, 3731–3736 (2004).

10. Chandran, V. I., Mansson, A. S., Barbachowska, M., Cerezo-Magana, M., Nodin, B., Joshi, B., KoppaD.A., N., Saad, O. M., Gluz, O., Isaksson, K., Borgquist, S., Jirstrom, K., Nabi, I. R., Jernstrom, H. & Belting, M. Hypoxia attenuates trastuzumab uptake and trastuzumab-emtansine (T-DM1) cytotoxicity through redistribution of phosphorylated caveolin-1. Molecular Cancer Research 18, 644–656 (Apr. 2020).

11. Osada-Oka, M., Kuwamura, H., Imamiya, R., Kobayashi, K., Minamiyama, Y., Takahashi, K., Tanaka, M. & Shiota, M. Suppression of the doxorubicin response by hypoxia-inducible factor-1α is strictly dependent on oxygen concentrations under hypoxic conditions. European Journal of Pharmacology 920 (Apr. 2022).

12. Martin, J. D., Cabral, H., Stylianopoulos, T. & Jain, R. K. Improving cancer immunotherapy using nanomedicines: progress, opportunities and challenges. Nature Reviews Clinical Oncology 17, 251–266 (Apr. 2020).

13. Rebucci, M. & Michiels, C. Molecular aspects of cancer cell resistance to chemotherapy May 2013.

14. Goldman, D. Theoretical models of microvascular oxygen transport to tissue. Microcirculation 15, 795–811 (2008).

15. Secomb, T. W. Blood Flow in the Microcirculation. Annual Review of Fluid Mechanics 49, 443–461 (2017).

16. Jain, R. K., Martin, J. D. & Stylianopoulos, T. The Role of Mechanical Forces in Tumor Growth and Therapy. Annual Review of Biomedical Engineering 16, 321–346 (2014).

17. Bernabeu, M. O., Köry, J., Grogan, J. A., Markelc, B., Beardo, A., D’Avezac, M., Enjalbert, R., Kaeppler, J., Daly, N., Hetherington, J., Krüger, T., Maini, P. K., Pitt-Francis, J. M., Muschel, R. J., Alarcón, T. & Byrne, H. M. Abnormal morphology biases haematocrit distribution in tumour vasculature and contributes to heterogeneity in tissue oxygenation. Proceedings of the National Academy of Sciences 117, 27811–27819 (2020).

18. Enjalbert, R., Krüger, T. & Bernabeu, M. O. Effect of vessel compression on blood flow in microvascular networks and its implications for tumour tissue hypoxia. Communications Physics 7 (Feb. 2024).

19. Enjalbert, R., Hardman, D., Krüger, T. & Bernabeu, M. O. Compressed vessels bias red blood cell partitioning at bifurcations in a hematocrit-dependent manner: Implications in tumor blood flow. Proceedings of the National Academy of Sciences 118 (2021).

20. Merlo, A., Berg, M., Duru, P., Risso, F., Davit, Y. & Lorthois, S. A few upstream bifurcations drive the spatial distribution of red blood cells in model microfluidic networks. Soft Matter 18, 1463–1478 (Feb. 2022).

21. Martin, J. D., Lanning, R. M., Chauhan, V. P., Martin, M. R., Mousa, A. S., Kamoun, W. S., Han, H. S., Lee, H., Stylianopoulos, T., Bawendi, M. G., Duda, D. G., Brown, E. B., Padera, T. P., Fukumura, D. & Jain, R. K. Multiphoton Phosphorescence Quenching Microscopy Reveals Kinetics of Tumor Oxygenation during Antiangiogenesis and Angiotensin Signaling Inhibition. Clinical Cancer Research 28, 3076–3090 (July 2022).

22. Hill, S. A., Pigott, K. H., Saunders, M. I., Powell, M. E. B., Arnold, S., Obeid, A., Ward, G., Leahy, M., Hoskin, P. J. & Chaplin, D. J. Microregional blood flow in murine and human tumours assessed using laser Doppler microprobes. British Journal of Cancer 74, 260–263 (1996).

23. Kimura, H., Braun, R., Ong, E., Hsu, R., Secomb, T., Papahadjopoulos, D., Hong, K. & Dewhirst, M. Flucuations in red cell flux in tumor microvessels can lead to transient hypoxia and reoxygenation in tumor parenchyma. Cancer Research 56, 5522–5528 (1996).

24. Michiels, C., Tellier, C. & Feron, O. Cycling hypoxia: A key feature of the tumor microenvironment. Biochimica et Biophysica Acta - Reviews on Cancer 1866, 76–86 (2016).

25. Suwa, T., Kobayashi, M., Nam, J. M. & Harada, H. Tumor microenvironment and radioresistance June 2021.

26. Grimes, D. R., Fletcher, A. G. & Partridge, M. Oxygen consumption dynamics in steady-state tumour models. Journal of the Royal Society, Interface / the Royal Society 1 (2014).

27. Fang, L., He, Y., Tong, Y., Hu, L., Xin, W., Liu, Y., Zhong, L., Zhang, Y. & Huang, P. Flattened microvessel independently predicts poor prognosis of patients with non-small cell lung cancer. Oncotarget 8, 30092–30099 (2017).

28. Fredrich, T., Welter, M. & Rieger, H. Tumorcode: A framework to simulate vascularized tumors. European Physical Journal E 41 (Apr. 2018).

29. Rieger, H., Fredrich, T. & Welter, M. Physics of the tumor vasculature: Theory and experiment. European Physical Journal Plus 131, 1–24 (Feb. 2016).

30. Pries, A., Ley, K., Classen, M. & Gaehtgens, P. Red Cell Distribution at Microvascular. Microvascular research 38, 81–101 (1989).

31. Pries, A. R., Neuhaus, D. & Gaehtgens, P. Blood viscosity in tube flow: Dependence on diameter and hematocrit. American Journal of Physiology - Heart and Circulatory Physiology 263 (1992).

32. Pries, A. R. & Secomb, T. W. Microvascular blood viscosity in vivo and the endothelial surface layer. American Journal of Physiology - Heart and Circulatory Physiology 289, 2657–2664 (2005).

33. Pries, A. R. & Secomb, T. W. Blood Flow in Microvascular Networks. In: Comprehensive Physiology. Handbook of Physiology: Microcirculation, 1–34 (2008).

34. Grogan, J. A., Connor, A. J., Markelc, B., Muschel, R. J., Maini, P. K., Byrne, H. M. & Pitt-Francis, J. M. Microvessel chaste: an open library for spatial modeling of vascularized tissues. Biophysical Journal 112, 1767–1772 (2017).

35. Alarcón, T., Byrne, H. M. & Maini, P. K. A cellular automaton model for tumour growth in inhomogeneous environment. Journal of theoretical biology 225, 257–274 (2003).

36. Owen, M. R., Stamper, I. J., Muthana, M., Richardson, G. W., Dobson, J., Lewis, C. E. & Byrne, H. M. Mathematical modeling predicts synergistic antitumor effects of combining a macrophage-based, hypoxia-targeted gene therapy with chemotherapy. Cancer Research 71, 2826–2837 (Apr. 2011).

37. Girish, S., Gupta, M., Wang, B., Lu, D., Krop, I. E., Vogel, C. L., Burris, H. A., LoRusso, P. M., Yi, J. H., Saad, O., Tong, B., Chu, Y. W., Holden, S. & Joshi, A. Clinical pharmacology of trastuzumab emtansine (T-DM1): An antibody-drug conjugate in development for the treatment of HER2-positive cancer. Cancer Chemotherapy and Pharmacology 69, 1229–1240 (May 2012).

38. McKeown, S. R. Defining normoxia, physoxia and hypoxia in tumours - Implications for treatment response. British Journal of Radiology 87 (Mar. 2014).

39. Fall, J., Ciannelli, L., Skaret, G. & Johannesen, E. Seasonal dynamics of spatial distributions and overlap between Northeast Arctic cod (Gadus morhua) and capelin (Mallotus villosus) in the Barents Sea. PLoS ONE 13 (Oct. 2018).

40. Anselin, L., Syabri, S. & Smirnov, O. Visualizing Multivariate Spatial Correlation with Dynamically Linked Windows. New Tools for Spatial Data Analysis: Proceedings of the Specialist Meeting (2002).

41. Padera, T., Stoll, B., Tooredman, J., Capen, D., di Tomaso, E. & Jain, R. Cancer cells compress intratumour vessels. Nature 427, 695 (2004).

42. Penta, R. & Ambrosi, D. The role of the microvascular tortuosity in tumor transport phenomena. Journal of Theoretical Biology 364, 80–97 (Jan. 2015).

43. Azzi, S., Hebda, J. K. & Gavard, J. Vascular permeability and drug delivery in cancers. Frontiers in Oncology 3 AUG (2013).

44. Chauhan, V. P., Martin, J. D., Liu, H., Lacorre, D. A., Jain, S. R., Kozin, S. V., Stylianopoulos, T., Mousa, A. S., Han, X., Adstamongkonkul, P., Popovíc, Z., Huang, P., Bawendi, M. G., Boucher, Y. & Jain, R. K. Angiotensin inhibition enhances drug delivery and potentiates chemotherapy by decompressing tumour blood vessels. Nature Communications 4 (2013).

45. Jain, R. K. Normalization of Tumor Vasculature: An Emerging Concept in Antiangiogenic Therapy. Science 307, 58–62 (2005).

46. Jain, R. K. Normalizing tumor microenvironment to treat cancer: Bench to bedside to biomarkers. Journal of Clinical Oncology 31, 2205–2218 (2013).

47. Pries, A. R., Secomb, T. W., Gaehtgens, P. & Gross, J. F. Blood flow in microvascular networks. Experiments and simulation. Circulation research, 826–834 (1990).

48. Lorthois, S., Cassot, F. & Lauwers, F. Simulation study of brain blood flow regulation by intra-cortical arterioles in an anatomically accurate large human vascular network: Part I: Methodology and baseline flow. NeuroImage 54, 1031–1042 (Jan. 2011).

49. Poon, K. A., Flagella, K., Beyer, J., Tibbitts, J., Kaur, S., Saad, O., Yi, J.-H., Girish, S., Dybdal, N. & Reynolds, T. Preclinical safety profile of trastuzumab emtansine (T-DM1): mechanism of action of its cytotoxic component retained with improved tolerability. Toxicology and applied pharmacology 273, 298–313 (2013).

50. Bordeau, B. M., Abuqayyas, L., Nguyen, T. D., Chen, P. & Balthasar, J. P. Development and Evaluation of Competitive Inhibitors of Trastuzumab-HER2 Binding to Bypass the Binding-Site Barrier. Frontiers in Pharmacology 13 (Feb. 2022).

